# Structure of the SMYD2-PARP1 Complex Reveals Both Productive and Allosteric Modes of Peptide Binding

**DOI:** 10.1101/2024.12.03.626679

**Authors:** Yingxue Zhang, Eid Alshammari, Jacob Sobota, Nicolas Spellmon, Emerson Perry, Tianxin Cao, Thamarahansi Mugunamalwaththa, Sheila Smith, Joseph Brunzelle, Gensheng Wu, Timothy Stemmler, Jianping Jin, Chunying Li, Zhe Yang

**Author notes:** Corresponding author: Zhe Yang, PhD, 540 East Canfield Street, Detroit, Michigan 48201, USA.

## Abstract

Allosteric regulation allows proteins to dynamically respond to environmental cues by modulating activity at sites away from the catalytic center. Despite its importance, the SET-domain protein lysine methyltransferase superfamily has been understudied. Here, we present four crystal structures of SMYD2, a unique family member with a MYND domain. Our findings reveal a novel allosteric binding site with high conformational plasticity and promiscuity, capable of binding peptides, proteins, PEG, and small molecules. This site exhibits positive cooperativity with substrate binding, influencing catalytic activity. Mutations here significantly alter substrate affinity, changing the enzyme’s kinetic profile. Specificity studies show interaction with PARP1 but not histones, suggesting targeted regulation. Interestingly, this site’s function remains unaffected by active site changes, indicating unidirectional mechanisms. Our discovery provides novel insights into SMYD2’s biochemical regulation and lays the foundation for broader research on allosteric control in lysine methyltransferases. Given SMYD2’s role in various cancers, this work opens exciting avenues for designing specific allosteric inhibitors with reduced off-target effects.

## Introduction

Allosteric regulation is a fundamental mechanism in biology that allows for rapid adaptation to environmental changes by directly regulating protein activity (*1*). This intrinsic property of proteins enables binding at one site to significantly influence activity at another. In cellular signaling, allosteric regulation is essential for cells to respond precisely to different signals through complex allosteric interactions (*2*). In metabolism, it plays a pivotal role by allowing cells to dynamically control metabolic flux in response to changing demands through feedback allosteric inhibition (*3*). Despite significant advances, studying allosteric regulation remains challenging due to the inherent complexity of allostery. One important class of proteins, the SET-domain protein lysine methyltransferase superfamily, which is crucial for phenotypic adaptation and epigenetic regulation, has significantly lagged behind other protein families, such as protein kinases, in allosteric studies (*4*). A critical barrier hampering these studies has been the lack of identification of any functional allosteric sites to which effector molecules can bind. Another significant reason is that current research on protein lysine methyltransferases has mainly focused on their downstream effects: through methylation of histone proteins, they are critical for governing the complex spatiotemporal patterns of gene expression during cell differentiation (*5*), and through methylation of non-histone targets, they modulate key signaling pathways involved in cell cycle control and stress responses (*6*). However, the absence of knowledge about allosteric sites in this superfamily limits our understanding of how environmental factors influence these proteins and how they translate these influences into responses that alter signaling pathways and epigenetic processes. Identifying allosteric sites could fundamentally enhance our understanding of the molecular mechanisms governing this enzyme superfamily.

Here, we present the discovery of a promiscuous allosteric site in SMYD2, a unique member of the protein lysine methyltransferase family characterized by the co-presence of the SET and MYND domains (*7*). We demonstrate that this new binding site confers positive cooperativity with substrate binding at the active site, capable of binding peptides, proteins, PEG, and small molecules. SMYD2 is a critical regulator of numerous cellular processes, methylating both histone and non-histone proteins to control chromatin remodeling, DNA damage response, cell cycle progression, cell differentiation, cell survival, and inflammation (*8, 9*). For instance, SMYD2 mono-methylates PARP1 at lysine 528, enhancing poly(ADP-ribosyl)ation activity and promoting poly(ADP-ribose) formation under oxidative stress (*10*). Beyond PARP1, more than ten other proteins have been individually characterized as SMYD2 targets, including histone H3, histone H4, p53, ERα, retinoblastoma protein 1, HSP90, β-catenin, STAT3, and the p65 subunit of NF-κB (*10*). Furthermore, over 150 additional proteins in the human genome have been identified as SMYD2 substrates *in vitro* (*11*). This broad substrate specificity raises intriguing questions about how the substrate selectivity of SMYD2 is spatiotemporally controlled and the biochemical mechanisms governing its activity, both of which remain largely unexplored.

Understanding these mechanisms is not only fundamentally significant but also of therapeutic interest. Most SMYD2 targets are tumor suppressors and oncogenes (*12*). This aligns with the fact that SMYD2 is overexpressed in various cancers and associated with poor patient outcomes (*13*). As a result, SMYD2 has become a focal point for drug design, with significant efforts directed at developing inhibitors targeting SMYD2 (*14–17*). While these inhibitors have shown promise in inhibiting cancer growth in both cell lines and mouse models, significant off-target effects have also been observed (*8, 18*). This issue is mainly because current SMYD2 inhibitors target only the highly conserved active site among SET-domain methyltransferases, increasing the likelihood of off-target effects. Our identification of a novel allosteric site in SMYD2 provides new insights into protein regulation. This discovery not only reveals a new biochemical mechanism regulating SMYD2 but also suggests potential target sites for future drug design. From a broader perspective, this finding could lay the groundwork for a better understanding of allosteric control mechanisms in the SET-domain protein lysine methyltransferase superfamily.

## Results

### 1. A Promiscuous Secondary Binding Site

We have determined four crystal structures of SMYD2, including a wild-type protein structure in complex with a PARP1 peptide, a wild-type protein structure in complex with an ethylene glycol polymer (PEG), and two structures of SMYD2 mutants (Fig. 1, 2A, 2B & S1). These structures reveal a novel promiscuous secondary binding site in SMYD2, which, together with the substrate binding site, binds to two separate PARP1 peptides (Fig. 1A & 1B). In terms of the overall fold, there are no significant differences between these structures (Fig. 2B). In all structures, SMYD2 maintains a bilobal fold, with the N- and C-lobes displaying a similar orientation and a similar degree of opening. The arrangement of the individual domains is also well maintained, with the SET domain located within the N-lobe, surrounded by the MYND, SET-I, and post-SET domains, and with the CTD domain forming the C-lobe (Fig. 1A & 2B). The structures are also well-preserved at the cofactor binding site, the substrate binding site, and the target lysine access channel (Fig. 2C-E).

**Fig. 1.**
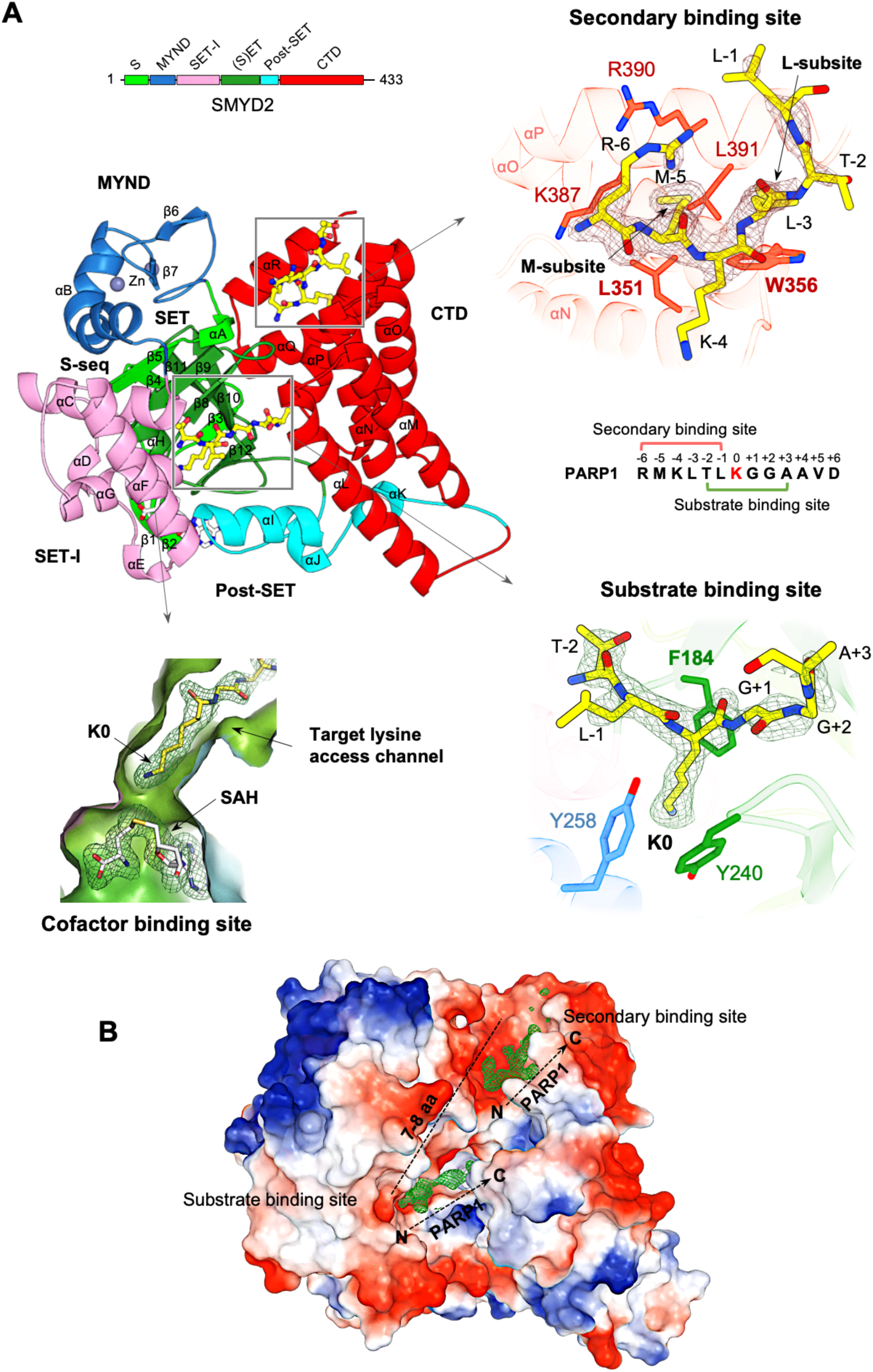
A novel secondary binding site. (A) Ribbon representation of the SMYD2-PAPR1 complex structure. The S-sequence, MYND, SET-I, core SET, post-SET, and CTD domains are depicted in light green, blue, pink, green, cyan, and red, respectively. Secondary structures, including α-helices and β-strands, are labeled and numbered according to their position in the sequence. AdoHcy and PARP1 peptides are shown as sticks with carbon atoms colored in white and yellow, respectively. Zinc atoms are depicted as spheres. The secondary binding site, substrate binding site, target lysine access channel, and cofactor binding site are indicated. The regions bound by SMYD2 are mapped onto the PARP1 peptide sequence. A schematic diagram of SMYD2 domains is colored according to the same scheme as in the structure. *2F_O_ - F_c_* omit maps of AdoHcy and PARP1 peptides were calculated at 2.1 Å and contoured at 1.0 σ. (B) SMYD2-PARP1 structure represented by molecular surface. The coloring corresponds to the electrostatic potential of the surface: red, white, and blue correspond to negative, neutral, and positive potential, respectively. PARP1 peptides are represented by their electron densities. The N- and C-termini of the peptides are labeled. The approximate number of amino acids that could link the two peptides is indicated.

**Fig. 2.**
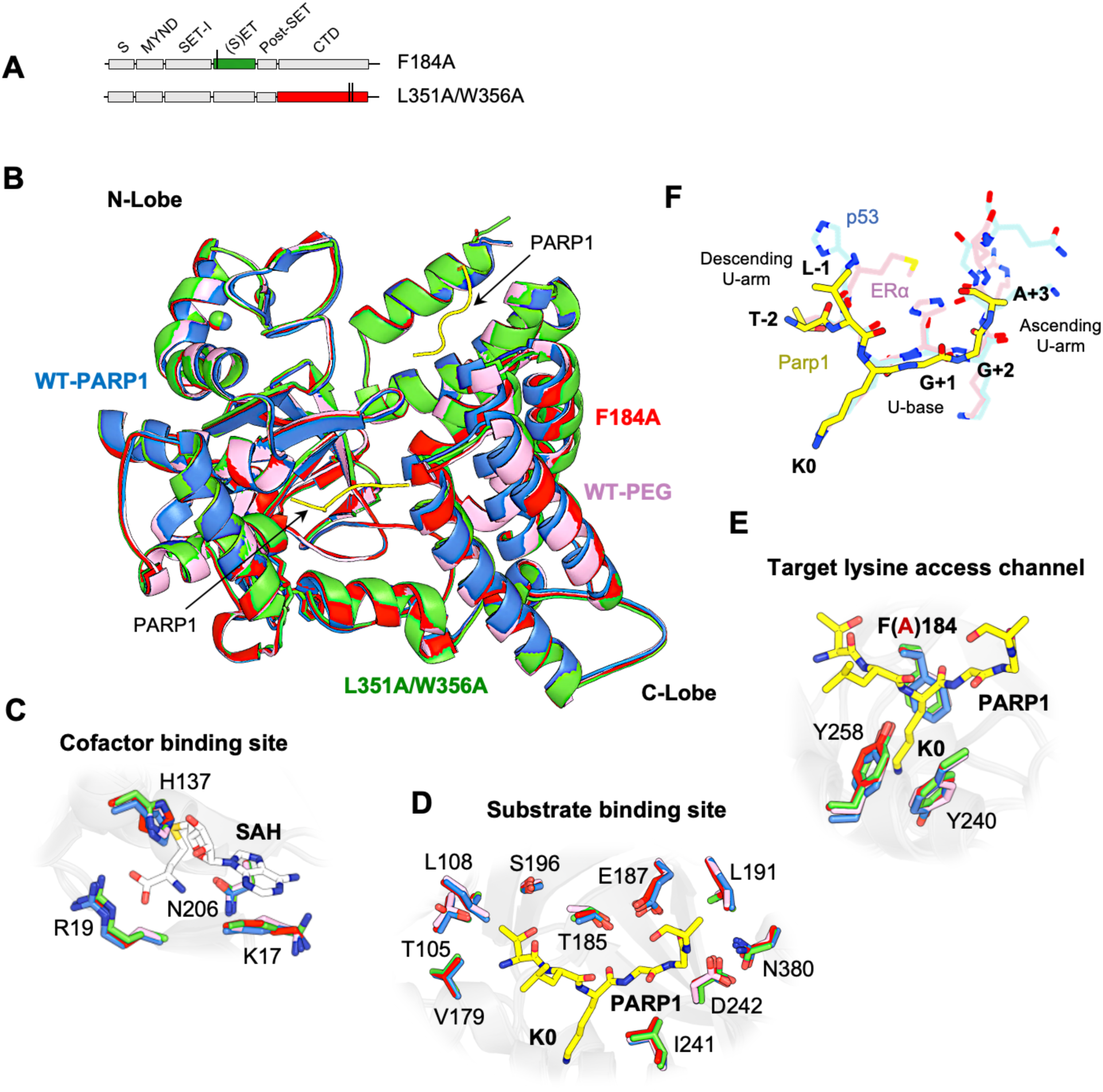
Structures of peptide-binding site mutants. (A) Constructs of SMYD2 mutants. (B) Structural superposition of SMYD2-PARP1 (blue), SMYD2-F184A (red), SMYD2-L351A/W356A (green), and SMYD2-PEG (pink) for the overall structure, (C) at the cofactor binding site, (D) at the substrate binding site, and (E) at the target lysine access channel. AdoHcy and PARP1 peptides are depicted as sticks with carbon atoms colored in white and yellow, respectively. (F) Structural superposition of SMYD2 substrates bound at the substrate binding site: PARP1 (yellow), p53 (blue), and Erα (pink).

Two peptide binding sites are identified in the wild-type SMYD2 structure in complex with a Lys528-containing PARP1 peptide (Fig. 1A & 1B). One corresponds to the well-known substrate binding site, and the other is new and had not been characterized before. At the substrate binding site, the PARP1 peptide binds in a U-shaped conformation, sandwiched by the β8–β9 hairpin from the SET domain and a loop connecting the SET and post-SET domains (Fig. 1A & 2F). The target lysine (Lys528) is located at the U-base of the peptide, with its side chain inserted into the target lysine access channel. In this position, the epsilon nitrogen of Lys528 is adjacent to AdoHcy at the cofactor binding site. AdoHcy is the byproduct formed following the methyl transfer from AdoMet (Fig. 1A & S1). This transfer reaction occurs at the junction between the cofactor binding site and the target lysine access channel.

The novel secondary binding site is located near the substrate binding site in the cleft between the N- and C-lobes (Fig. 1A & 1B). Three regions contribute to forming this site: four α-helices from the CTD domain, the turn region of the β8–β9 hairpin, and an α-helix and its following loop from the MYND domain. This site is mostly hydrophobic, composed of eight hydrophobic and two hydrophilic amino acids. Two subsites were defined based on their interactions with the peptide: the M-subsite interacts with Met-5 of the peptide, while the L-subsite interacts with Leu-3. Key residues for peptide binding at the M-subsite include Leu351, Lys387, Arg390, and Leu391, which form a shallow hydrophobic pocket where Met-5 is bound. At the L-subsite, Trp356 is important for peptide binding, as Leu-3 is stacked between the side chain of this residue and the main chains of Arg390 and Gly394.

Due to their proximity, it might seem possible for the secondary binding site and the substrate binding site to bind the same PARP1 peptide. However, structural analysis does not support this. The secondary binding site binds the N-terminal region of the peptide, while the substrate binding site binds the middle region around the target lysine (Fig. 1A). Their binding regions overlap by two residues, making it impossible for them to bind the same PARP1 peptide simultaneously. Additionally, the orientations of the two peptides do not support this possibility. For them to bind the same peptide, the C-terminal end of the peptide at the secondary binding site would need to be sufficiently close to the N-terminal end of the peptide at the substrate binding site (Fig. 1B). However, these two ends are over 25 Å apart in the structure. Based on this analysis, we conclude that SMYD2 binds to two individual PARP1 peptides in the structure.

The secondary binding site is promiscuous, capable of binding to other molecules. The polymer (PEG) (Fig. 3A). This finding was also unexpected since PEG was primarily used as a precipitant in crystallization experiments. In this structure, PEG binds across the L and M subsites and extends into an additional binding site in the CTD domain. At the secondary binding site, PEG mimics the binding of the PAPR1 peptide, with the polymer chain running parallel to the side chains of Leu-3 and Met-5. At the additional binding site, PEG binds to a groove formed by the first three α-helices of the CTD domain. Though unintentional, this structure shows that the secondary binding site can bind a PEG polymer, demonstrating its binding promiscuity.

**Fig. 3.**
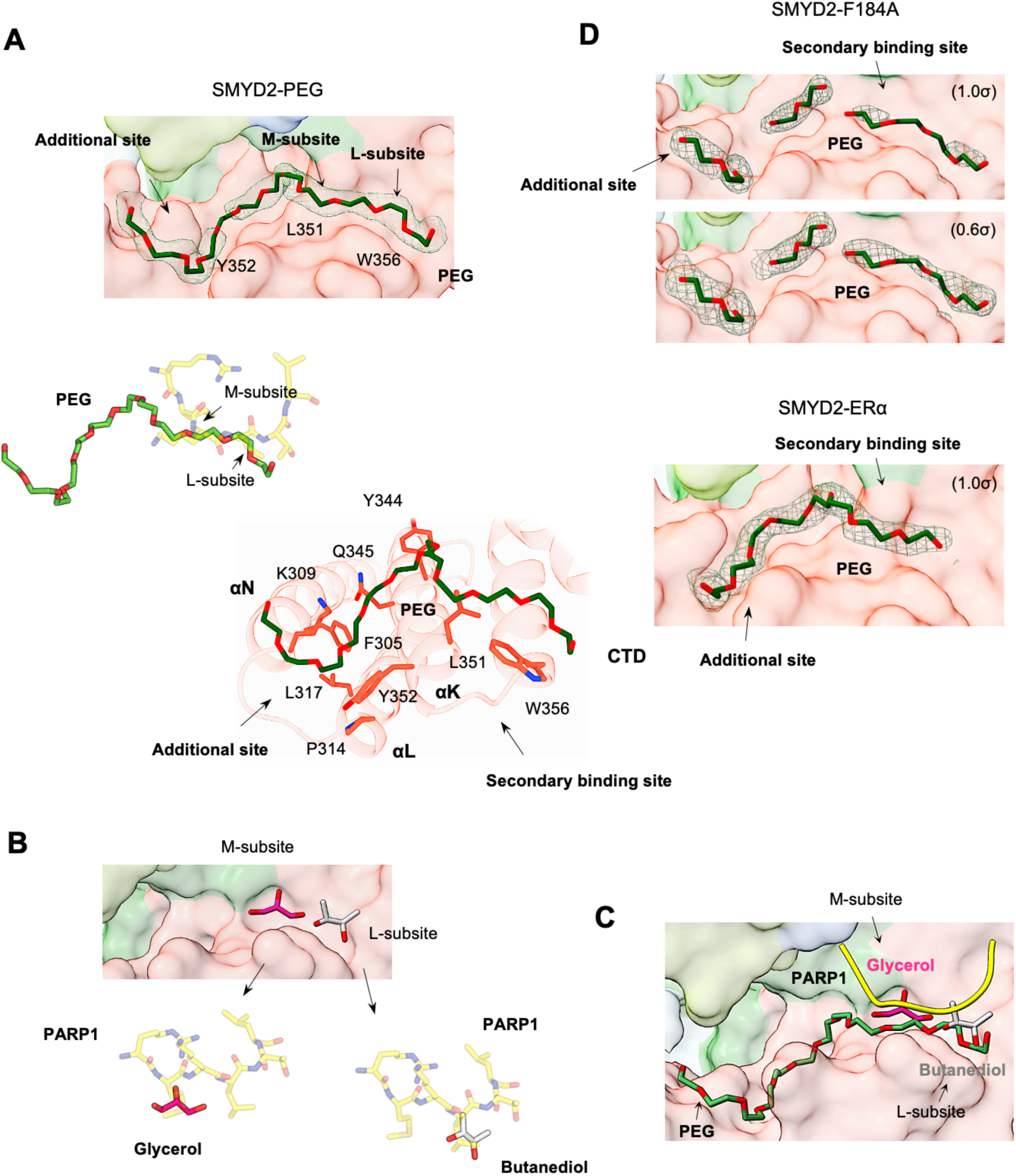
Promiscuous secondary binding site. (A) The PEG binding site in the SMYD2-PEG structure. Surface representation of the PEG binding site (top), superposition of PEG and the PARP1 peptide (middle), and SMYD2 residues involved in PEG binding (bottom). PEG, the PARP1 peptide, and SMYD2 residues are depicted as sticks with carbon atoms colored in dark green, yellow, and red, respectively. Residues important for forming the secondary binding site, including L35l, Y352, and W356, are labeled on the surface. (B) Butanediol (grey) and glycerol (magenta) bound at the secondary binding site and their superposition with the PARP1 peptide. (C) Promiscuous secondary binding site overlaid with the PARP1 peptide (yellow), PEG (dark green), butanediol (grey), and glycerol (magenta). (D) The PEG binding site in the SMYD2-F184A and SMYD2-ERα structures. 2F_O_ - *F_c_* omit maps of PEG molecules were contoured at 1.0 σ for SMYD2-PEG and SMYD2-ERα, and 1.0 σ and 0.6 σ for SMYD2-F184A.

In addition to peptides and PEG, the secondary binding site can also bind small molecules (Fig. 3B). A survey of all SMYD2 structures in the Protein Data Bank (PDB) (24 structures) identifies two where the secondary binding site is bound by small molecules. In one structure (PDB code: 3S7D), a butanediol is found at subsite L, while in the other structure (PDB code: 3TG4), subsite M is bound by a glycerol. The butanediol superimposes well with the Leu-3 side chain of the PARP1 peptide, while the glycerol overlays with the Met-5 side chain. These findings not only further demonstrate the promiscuity of the secondary binding site but also confirm the importance of both subsites M and L in molecular binding. Nonetheless, these SMYD2 structures show that the secondary binding site is inherently promiscuous and can bind to peptides, PEG, and small molecules (Fig. 3C).

### 2. A Secondary Binding Site Mutation Disrupts Substrate Binding

To explore the function of the secondary binding site and its connection to the substrate binding site, we generated two SMYD2 mutants with specific point mutations within these sites. L351A/W356A is a secondary-binding-site mutation that targets both subsites M and L, while F184A targets the target lysine access channel at the active site (Fig. 2A). We determined the crystal structures of these mutants to reveal any changes in peptide binding at each site (Fig. S1). In the L351A/W356A mutant structure, neither the secondary binding site nor the substrate binding site contains a peptide, although the PARP1 peptide was included during crystallization. This result suggests two possibilities: one is that this mutation disrupts binding at both sites, and the other is that the binding of the peptide to the substrate binding site depends on another peptide binding to the secondary binding site. Both scenarios suggest that the secondary binding site is crucial in regulating SMYD2 substrate binding, potentially by allosterically modulating the structure to facilitate effective substrate engagement. It is important to note that the experimental conditions for crystallizing both the wild-type and mutant proteins were identical, and their crystals grew in the same space group (Table S2). Despite different unit cell dimensions, the crystal packing was similar at both the substrate binding site and the secondary binding site (Fig. S2). These similarities rule out the possibility that the experimental conditions and crystal packing caused the observed differences in peptide binding.

Our initial hypothesis was that the L351A/W356A mutation would cause significant structural changes, disrupting both binding sites. However, this only explains the secondary binding site, as the substrate binding site shows no significant structural changes (Fig. 2D). Comparing the wild-type and mutant structures reveals that the mutation disrupts the integrity of the secondary binding site, making it less defined, explaining the loss of binding there (Fig. 4A). Several minor structural adjustments are associated with this mutation. The most notable one is a slight movement of helices L, M, N, and O relative to helix P. This movement is likely allowed due to the extra space created by the mutation. Another difference is a slight shift of a hydrophobic core in the same direction. These findings suggest that the L351A/W356A mutation affects the secondary binding site but does not cause widespread structural disruption beyond the local region of the mutation site. Consistently, the substrate binding site, located about 8 Å from the mutation site, remains largely unaffected by the mutation. Key residues involved in substrate binding are well-aligned between the wild-type and mutant structures (Fig. 2D & 2E). These residues include Ser196, Val179, Thr105, and Leu108, which recognize the Leu-1 of the peptide; Phe184, Tyr240, and Tyr258, which form a channel for the target lysine to bind; Glu187, Thr185, and Ile241, which stabilize the backbone of the peptide; and Leu191, Asn380, and Asp242, which stack against the C-terminus of the peptide. These findings indicate that the L351A/W356A mutation does not visibly alter the substrate binding site, making it unclear how this mutation affects substrate binding.

**Fig. 4.**
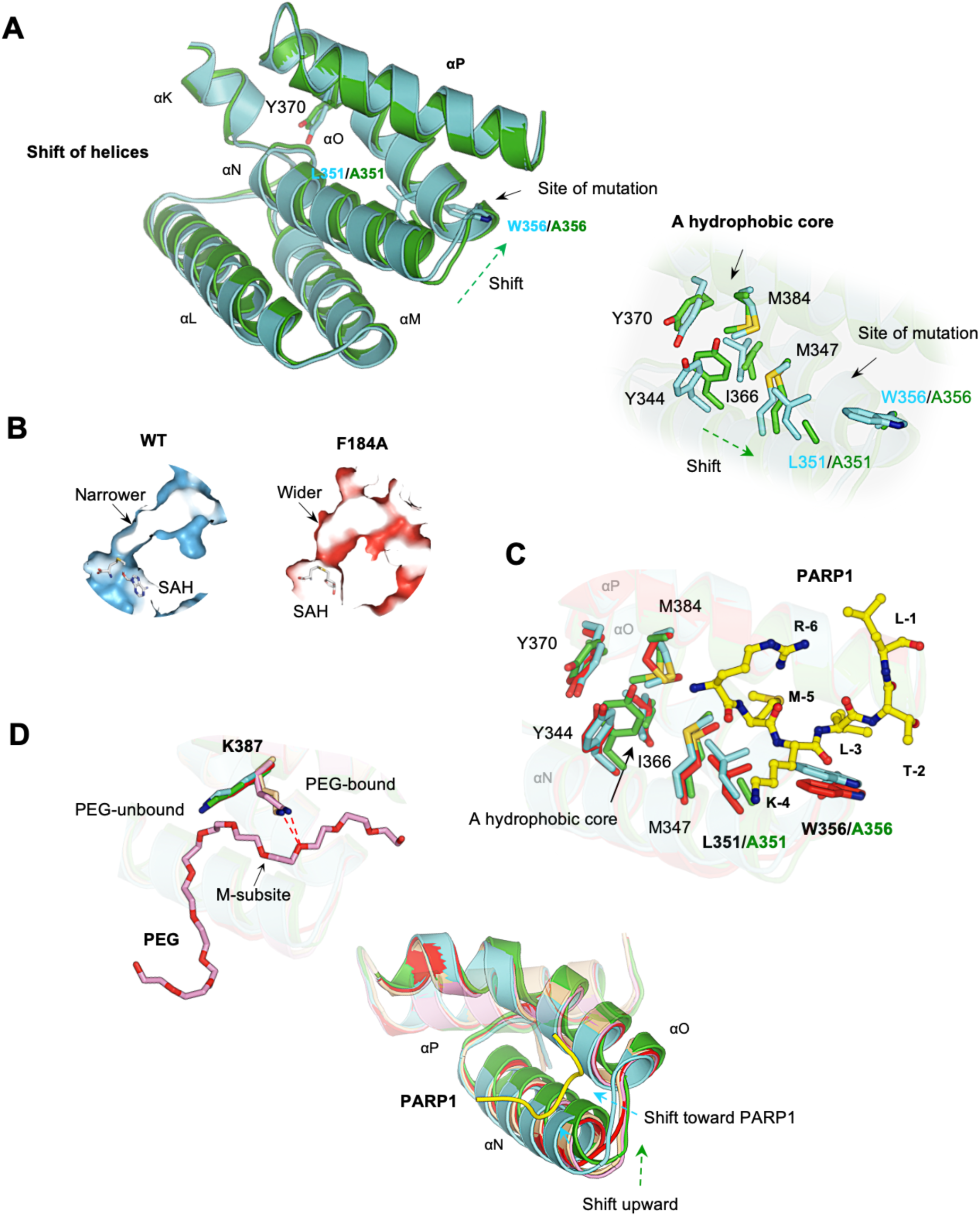
Structural comparison of wild-type and mutant SMYD2. (A) Minor structural differences at the secondary binding site between SMYD2-PARP1 (cyan) and SMYD2-L351A/W356A (green). Left: shift of helices. Right: shift of a hydrophobic core. (B) The widened target lysine access channel in SMYD2-F184A. (C) Superposition of SMYD2-PARP1 (cyan), SMYD2-F184 (red), and SMYD2-L351A/W356A (green) at the secondary binding site. (D) The structural flexibility of the secondary binding site. Left: PEG­bound and PEG-unbound structures have different K387 conformations. Right: the unique structural changes upon PARP1 binding, PEG binding, and point mutation.

### 3. An Active-Site Mutation Does Not Affect PEG Binding at the Secondary Binding Site

The F184A mutant was also crystallized with the PARP1 peptide, but no peptide is found in either the substrate binding site or the secondary binding site (Fig. S1), suggesting that this mutation also disrupts binding at both sites. The loss of binding at the substrate binding site can be readily explained by the structure. Phe184, together with the two other aromatic residues Tyr240 and Tyr258, forms the target lysine access channel (Fig. 1A). Mutating Phe184 to the smaller alanine (F184A) widens this channel significantly (Fig. 4B). In the wild-type structure, the target lysine fits snugly within the channel (Fig. 1A). The mutation likely disrupts this precise fit, leading to the loss of binding. However, it is unclear how this mutation disrupts the secondary binding site. Compared to L351A/W356A, the F184A mutation causes even fewer structural changes. No significant adjustments are found around the mutation site, and the active site residues are highly superimposed between the wild-type and mutant structures (Fig. 2C-E). The secondary binding site, about 8 Å away from the mutation site, is not significantly affected either. The residues involved in peptide binding, including those forming subsites M and L, are well aligned (Fig. 4C), making it unclear why no peptide was found at the secondary binding site.

Despite no peptide being identified, the secondary binding site contains PEG-like electron densities (Fig. 3D). These densities occupy positions similar to where PEG is found in the SMYD2-PEG structure (Fig. 3A). They fill both subsites M and L, although not as continuous as in the SMYD2-PEG structure. This suggests that the F184A mutation does not significantly alter PEG binding at the secondary binding site. In a broader sense, this discovery implies that the binding ability of the secondary binding site is not affected by changes in the substrate binding site. Their interaction appears to be unidirectional, from the secondary binding site to the substrate binding site, but not vice versa. To support this idea, we analyzed all PEG-bound SMYD2 structures, including the SMYD2-ERα structure (*19*). At least for the PEG molecule, binding to the secondary binding site appears to be independent of the status of the substrate binding site. A PEG molecule can bind to the secondary binding site regardless of whether the substrate binding site is mutated (SMYD2-F184A structure), bound by an ERα peptide (SMYD2-ERα structure), or without peptide binding (SMYD2-PEG structure) (Fig. 3). This finding confirms that the substrate binding site does not communicate changes to the secondary binding site, showing that the secondary binding site remains functional independent of changes in the substrate binding site.

### 4. The Secondary Binding Site with High Conformational Plasticity

Comparison of five SMYD2 structures, including SMYD2-PARP1, SMYD2-L351A/W356A, SMYD2-F184A, SMYD2-PEG, and SMYD2-ERα, reveals that the secondary binding site has high conformational plasticity, being more sensitive to perturbations than the substrate binding site (Fig. 4D). The substrate binding site remains structurally stable regardless of peptide binding or mutations, with all key residues aligned across structures bound by PARP1 peptide, ERα peptide, with mutations, and without any peptide (Fig. 2D & 2E). In contrast, the secondary binding site shows distinct responses to different structural perturbations. In PEG-bound structures, the side chain of Lys387 adopts a bent conformation, pointing up to the PEG molecule and forming a hydrogen bond with an oxygen atom from PEG (Fig. 4D). Without PEG, this side chain assumes a stretched conformation, which contributes to forming the M subsite for peptide binding. Another specific change is found at helices N and O (Fig. 4D). When a peptide is bound, these two helices move slightly inward toward the bound peptide, resulting in a slightly narrower site than when PEG is bound. The location of these helices is also affected by the L351A/W356A mutation, but the direction of this change is different from that caused by peptide binding (Fig. 4D). These findings suggest that the secondary binding site is more susceptible to structural changes than the substrate binding site. This high conformational plasticity may be a key mechanism underlying its binding promiscuity, allowing it to adapt to and interact with structurally diverse molecules.

### 5. The Secondary Binding Site Exhibits Positive Cooperativity with Substrate Binding

To confirm that SMYD2 contains two peptide-binding sites, we analyzed the binding between SMYD2 and PARP1 using isothermal titration calorimetry (ITC) with the same PARP1 peptide used in the crystal structure analysis (Fig. 5A). The results showed that the peptide binds in a 2:1 ratio with the wild-type protein, which changes to 1:1 with the F184A mutation (Fig. 5B & 5C). This confirms that SMYD2 has two peptide-binding sites, indicating that the F184A mutation affects one site while leaving the other intact. This aligns with the F184A mutant structure, where the substrate binding site is empty but the secondary binding site can still bind PEG (Fig. S1). Strikingly, the L351A/W356A mutation completely abolished peptide binding, as evidenced by the absence of ITC heat changes (Fig. 5A & 5B). This indicates that this mutation disrupts both binding sites, consistent with the L351A/W356A mutant structure, where neither site binds to a peptide or PEG (Fig. S1). Therefore, both crystallographic and ITC studies suggest that there is positive cooperativity in peptide binding: binding to the secondary binding site facilitates binding to the substrate binding site. To further support this notion, the wild-type data were fitted with the binding polynomial (Fig. 5B & 5C). The binding polynomial is a general model-free methodology that provides information about whether binding cooperativity exists, whether the cooperativity is positive or negative, and the magnitude of the cooperative energy (*20*). The fitting yielded β1 = 6.3×10^4^ M^-1^, ΔH_1_ = −4.2 kJ/mol, β2 = 2.5 x10^9^ M^-2^, and ΔH_2_ = −7.2 kJ/mol. The calculated value for π2 is 2.5, suggesting positive cooperativity between the two sites.

**Fig. 5.**
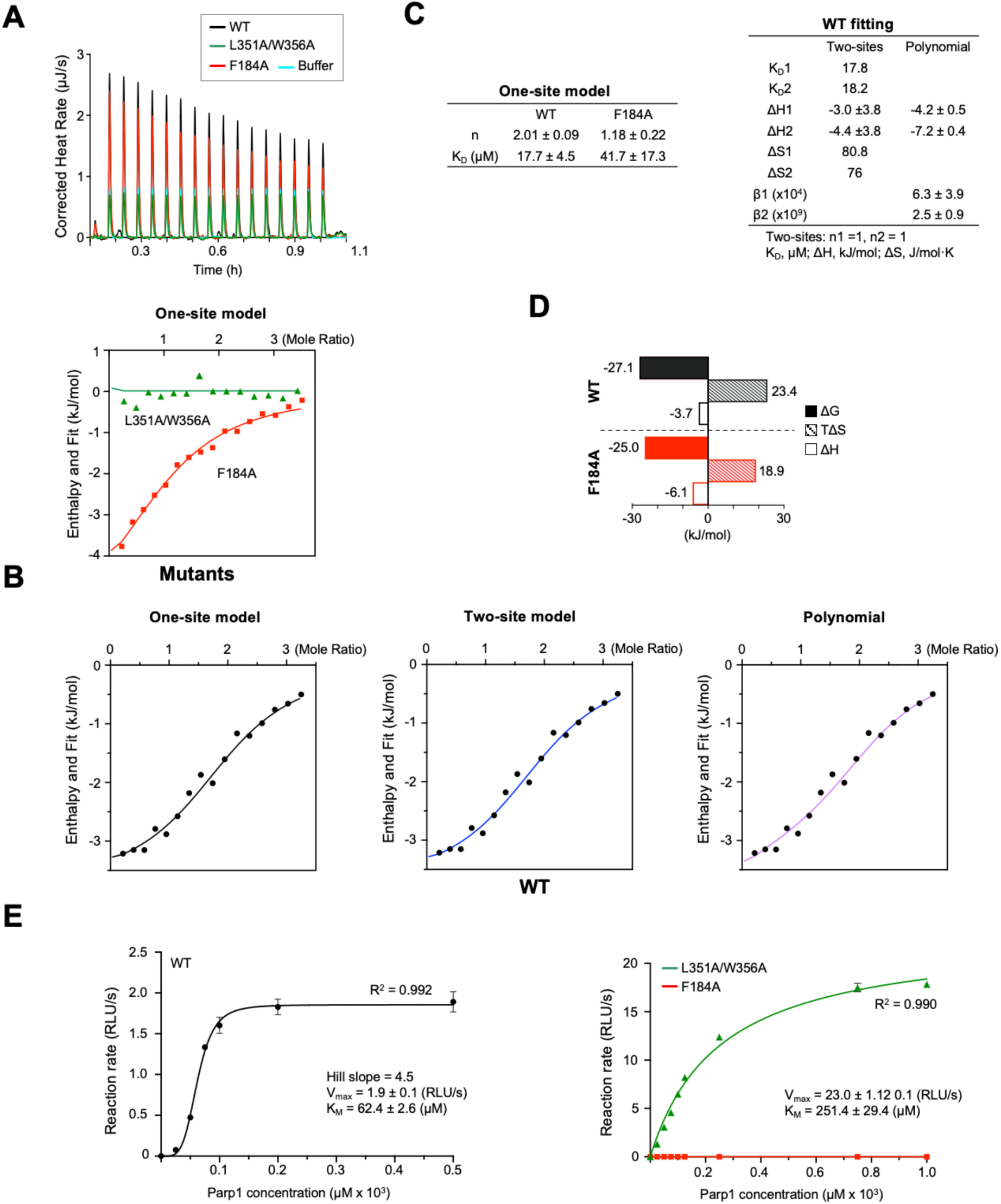
The secondary binding site exerts allosteric regulation. (A) Raw data ofITC analysis of the binding between SMYD2 proteins and the PARP1 peptide, including the control titration. (B) ITC binding isotherms with normalized heat changes. Top panel: Fl 84A and L351A/W356A data were fitted with the one-site model (red line) and the blank model (green line), respectively. Bottom panel: Wild-type SMYD2 data were fitted with the one-site (left), two-site (middle), and binding polynomial (right) models. (C) Thermodynamic parameters estimated from model fitting. (D) Entropic and enthalpic contributions to the binding free energy. (E) Steady-state enzyme kinetics of SMYD2 proteins with the PARPl peptide as the substrate. Each data point represents the average of three replicates; error bars represent the standard error. The luminescence readings, expressed as RLU (relative light unit), indicate enzymatic activities. Wild-type SMYD2 data were fitted to the Hill equation (black line). L351A/W356A and F184A data were fitted to the Michaelis-Menten equation (green and red lines). Enzyme kinetic parameters are shown.

To understand the mechanism of molecular recognition, we analyzed the thermodynamic properties of SMYD2-PARP1 interactions by fitting the ITC data to one-site and two-site models (Fig. 5B). Both models fit well with the wild-type data and showed similar dissociation constants (K_D_). In the one-site model, the overall binding free energy (βG) had positive contributions from both enthalpy and entropy, with entropy playing a larger role (Fig. 5D). This suggests that peptide binding in SMYD2 is mainly driven by hydrophobic interactions. Binding enthalpy and entropy provide insights into the types of interactions driving the binding process (*21*). Hydrogen bonds and electrostatic interactions typically result in negative changes in enthalpy, while hydrophobic interactions usually lead to a significant increase in entropy. However, it is important to note that the parameters from the wild-type data reflect contributions from both binding sites, representing an overall binding behavior rather than individual sites. In contrast, the F184A mutant, which has only one functional site, reveals the nature of the secondary binding site. The F184A data were fitted with the one-site model, and the estimated K_D_ is similar to that of the wild-type data (Fig. 5B & 5C). Compared to the wild-type, this mutant exhibits a more balanced thermodynamic signature, though entropy still interaction between the PARP1 peptide and the secondary binding site, where both hydrophobic contacts and hydrogen bonding are involved, with hydrophobic interactions being more dominant (Fig. 1A). This correlation between structural and thermodynamic properties provides deeper insights into the binding mechanisms at the secondary binding site, revealing how molecular conformation and energetic changes govern the binding process.

### 6. The Secondary Binding Site Exerts Positive Allosteric Regulation

To determine whether the secondary binding site regulates the methyltransferase activity of SMYD2, we studied its steady-state kinetics using the PARP1 peptide as a substrate. Wild-type SMYD2 exhibits a sigmoidal kinetic curve, indicating the presence of more than one peptide-binding sites in the protein (Fig. 5D). The Hill coefficient is 4.5, suggesting positive cooperativity, where other binding sites enhance substrate binding at the active site. Strikingly, the L351A/W356A mutation shifts the kinetic profile from sigmoidal to classic Michaelis-Menten kinetics, exhibiting single-site hyperbolic kinetic behavior (Fig. 5D). This shift signifies the loss of positive cooperativity with the mutation. This suggests that the intact secondary binding site is essential for the cooperative substrate binding observed in wild-type SMYD2. A detailed analysis of the kinetic parameters further supports the critical role of the secondary binding site in regulating SMYD2 kinetics and substrate binding. The L351A/W356A mutation results in a more than 10-fold increase in *V_max_*, while *K_M_*, which reflects apparent substrate binding affinity, shows a 4-fold increase. The catalytic efficiency, *k*_cat_/*K_M_*, increases by 2.5-fold with the mutation. The increased *K_M_* suggests that the substrate binding affinity is reduced by the mutation, whereas the increased catalytic efficiency indicates that the mutation increases the enzyme’s overall ability to convert substrate to product. These kinetic studies demonstrate that the secondary binding site functions as an allosteric site, exerting positive allosteric regulation, where the binding of a PAPR1 peptide to this site enhances the binding of another PARP1 peptide to the active site.

### 7. The Secondary Binding Site Interacts With PARP1 Protein But Not Histones

To further characterize the interaction between SMYD2 and PARP1, we used a PARP1 protein fragment composed of residues 518–1014. This fragment contains binding sites for both the substrate binding site and the secondary binding site of SMYD2, as well as the entire PARP1 catalytic domain (Fig. 6A). We aimed to investigate the interaction between SMYD2 and PARP1 in the context of a PARP1 protein. This interaction was analyzed using an ELISA assay, where PARP1 protein was coated onto wells, and bound SMYD2 was detected by a specific SMYD2 antibody (*22*). The results reveal that wild-type SMYD2, the L351A/W356A mutant, and the F184A mutant all bind to the PARP1 protein. All binding curves exhibit a hyperbolic shape, with no significant differences in their binding abilities (Fig. 6B). The concentrations required for 50% saturation are 0.060 μM for wild-type SMYD2, 0.053 μM for the L351A/W356A mutant, and 0.085 μM for the F184A mutant. At high SMYD2 concentrations, when the binding was approaching saturation, all three SMYD2 proteins showed similar binding levels, suggesting that they bind to the PARP1 protein with the same stoichiometry. Similar results were obtained in a reciprocal ELISA binding experiment, where SMYD2 was coated onto wells and bound PARP1 was detected by a specific PARP1 antibody (Fig. S3). A GST-pulldown assay using GST-tagged SMYD2 proteins also showed similar levels of PARP1 protein being pulled down by all three SMYD2 proteins (Fig. 6C). These results indicate that, unlike the PARP1 peptide, the ability of the PARP1 protein to bind to SMYD2 is not significantly affected by these SMYD2 mutations. For the F184A mutant, the intact secondary binding site may be responsible for retaining the binding of the PARP1 protein, while the L351A/W356A mutant retains binding likely due to the intact substrate binding site, despite its weakened ability to bind to the PARP1 peptide (Fig. 5).

**Fig. 6.**
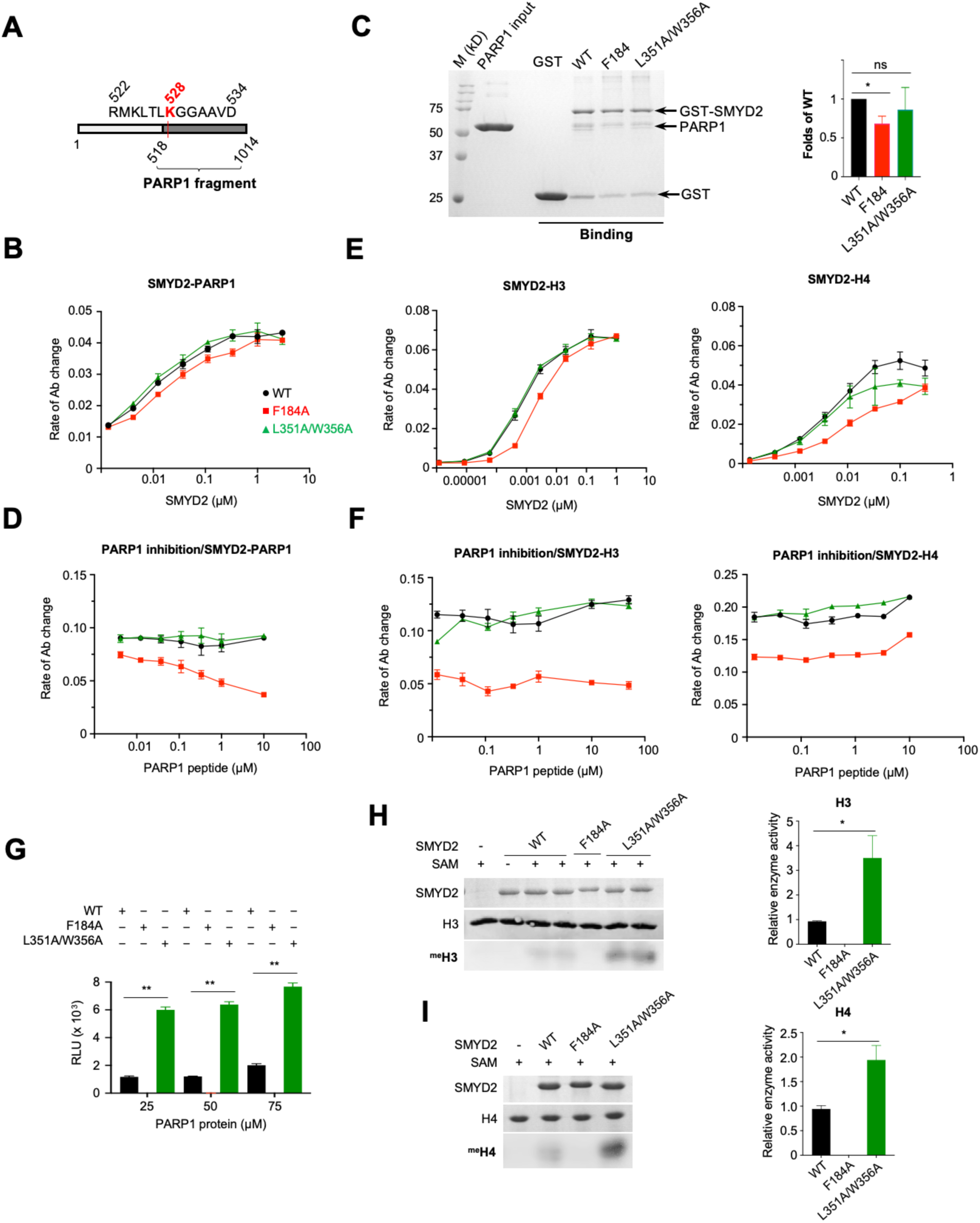
The binding specificity and regulatory nature of the secondary binding site. (A) The construct ofa PARPI protein fragment {PARP1{518-1014)). (B) ELISA and (C) GST pulldown analysis of the binding between SMYD2 proteins and PARP1{518-1014). (D) Inhibition ofSMYD2-PARP1{518-1014) interaction by the PARPl peptide. (E) ELISA analysis of the binding between SMYD2 proteins and histone H3 (left) and histone H4 (right). (F) Inhibition of SMYD2-H3 (left) and SMYD2-H4 (right) interactions by the PARPl peptide. (G) The methyltransferase activity ofSMYD2 proteins on PARP1(518-1014) assayed by the MTase-Glo methyltransferase assay. The methyltransferase activity of SMYD2 proteins on (H) H3 and (I) H4 assayed by the antibody-based methyltransferase assay. One(*) and two(**) indicate statistical significance with p-values :’.S 0.05 and 0.001, respectively.

To examine whether the secondary binding site is involved in the interaction with the PARP1 protein, we performed inhibition experiments using the PARP1 peptide as an inhibitor. SMYD2 proteins were pre-incubated with varying concentrations of the PARP1 peptide before assessing their binding to the coated PARP1 protein in an ELISA assay. The results reveal that the PARP1 peptide has no significant effect on the binding of wild-type SMYD2 or the L351A/W356A mutant. However, the binding of the F184A mutant is almost completely disrupted at high peptide concentrations (Fig. 6D). This suggests that, compared to wild-type SMYD2 or the L351A/W356A mutant, the F184A mutant is more sensitive to inhibition by the PARP1 peptide. The IC_50_ of this inhibition is approximately 0.153 μM, as estimated by fitting the dose-response curve with a four-parameter logistic equation. The ability of the PARP1 peptide to compete with the PARP1 protein for binding to the F184A mutant suggests that they may bind to the same site on the mutant. As shown by ITC experiments, the PARP1 peptide binds only at the secondary binding site on the F184A mutant (Fig. 5). This binding, combined with its ability to inhibit the binding of the PARP1 protein, suggests that this site may also be where the PARP1 protein binds. These findings collectively indicate that the secondary binding site is capable of binding its target site within the context of a PARP1 protein.

To evaluate the binding specificity of the secondary binding site, we examined the interactions between SMYD2 and histone H3 and between SMYD2 and histone H4 using an ELISA assay. In these experiments, H3 or H4 was coated onto wells, and bound SMYD2 was detected by a specific SMYD2 antibody. H3 and H4 are known substrates of SMYD2, binding at the substrate binding site (*7, 23*). We aimed to determine whether the secondary binding site is involved in their binding. All three SMYD2 proteins bound to H3 and H4 (Fig. 6E). The concentrations required for 50% saturation for wild-type SMYD2, the L351A/W356A mutant, and the F184A mutant are 0.060 μM, 0.023 μM, and 0.085 μM for H3, and 0.070 μM, 0.033 μM, and 0.095 μM for H4, respectively. The F184A mutation reduced binding to both H3 and H4, whereas the L351A/W356A mutation did not affect H3 binding but slightly reduced H4 binding. At high SMYD2 concentrations, when the binding was approaching saturation, all three SMYD2 proteins showed similar levels of binding to H3 or H4, suggesting that they bind to these histones with the same stoichiometry. To investigate the role of the secondary binding site in histone binding, we performed inhibition experiments using the PARP1 peptide as an inhibitor, similar to the approach used to study SMYD2 and PARP1 protein interactions. As observed with the PARP1 protein, the PARP1 peptide did not affect the binding of H3 or H4 to either wild-type SMYD2 or the L351A/W356A mutant at any tested concentration (Fig. 6F). However, unlike with the PARP1 protein, where the PARP1 peptide competed off the PARP1 protein from the F184A mutant, the binding of either histone to this mutant was unaffected by the PARP1 peptide. This suggests that the secondary binding site is not involved in the binding of histone H3 or H4.

### 8. The Secondary Binding Site Mutation Enhances SMYD2 Methyltransferase Activity

To further demonstrate the regulatory nature of the secondary binding site, we compared the methyltransferase activities of wild-type and mutant SMYD2 on various substrates. The L351A/W356A mutation enhances the methyltransferase activity of SMYD2 across all tested substrates, irrespective of its differential effects on substrate binding affinities. In the case of the PARP1 peptide, the *V_max_* of SMYD2 increases 10-fold with this mutation, despite a significant decrease in its apparent binding affinity to the active site (Fig. 5D). For the PARP1 protein, whose binding to SMYD2 is not affected by the mutation, there is also a notable increase in its methylation level (Fig. 6G). For H3 and H4, this mutation also results in a marked increase in their methylation, although it had varying effects on their binding to SMYD2 (Fig. 6H & 6I). These results indicate that the methyltransferase activity of SMYD2 is sensitive to perturbations at the secondary binding site, with the L351A/W356A mutation positively impacting the enzyme’s activity. In contrast, the F184A mutation at the target lysine access channel abolishes the methyltransferase activity of SMYD2 on all tested substrates, even though these substrates, except for the PARP1 peptide, can still bind to the mutant (Fig. 5D & 6). Together, these findings demonstrate that the target lysine access channel is essential for catalysis, whereas the secondary binding site plays a regulatory role.

## Discussion

How SMYD2 binding promiscuity and substrate specificity are temporally and spatially controlled remains largely unknown. Given the large number of potential substrates and the involvement of SMYD2 in diverse cellular pathways, its functional regulation is presumably complex, likely involving multiple lines of mechanisms. It has been shown that SMYD2 expression levels are temporally and spatially regulated during mouse heart development; it is preferentially expressed in the neonatal heart and less so in the embryonic and adult hearts (*24*). SMYD2 expression is also differentially regulated during cellular differentiation, being induced during the differentiation of human embryonic stem cells and preferentially expressed in somatic cells versus pluripotent cells (*25*). In this study, we identified a promiscuous allosteric site in SMYD2. The identification of this site provides new perspectives on understanding the functional regulation of SMYD2. This new binding site is regulatory in nature, not only being structurally sensitive to perturbations by molecular binding and point mutations but also capable of allosterically regulating substrate binding and catalytic activity. Like the substrate binding site, this allosteric site exhibits promiscuous binding properties, capable of binding peptides, proteins, an ethylene glycol polymer, and small molecules. These promiscuous binding properties may be attributable to the structural nature of this site, which is prone to subtle conformational changes upon ligand binding, thereby making it adaptable to accommodate various structurally different molecules.

From a broader perspective, identifying this allosteric site helps us understand allosteric control within the SET-domain protein lysine methyltransferase superfamily. This site might represent a general regulatory mechanism that evolved to control the SMYD protein family. For instance, SMYD1 and SMYD3 also have surface pockets in the same region as this secondary binding site (Fig. 7). These pockets are highly conserved across species and among SMYD proteins, suggesting that similar allosteric effects could occur in these SMYD proteins. In addition, the regulatory mechanism used by the secondary binding site may apply to other protein lysine methyltransferase families. Protein lysine methyltransferases have a complex structure with multiple domains, such as the pre-SET, SET-I, and post-SET domains, as well as subfamily-specific domains (*7*). This multi-domain structure could create new interacting sites with regulatory roles (*26*). Like SMYD2, allosteric regulation through these sites might be central to the role of these proteins in the epigenetic regulation of human development and disease.

**Fig. 7.**
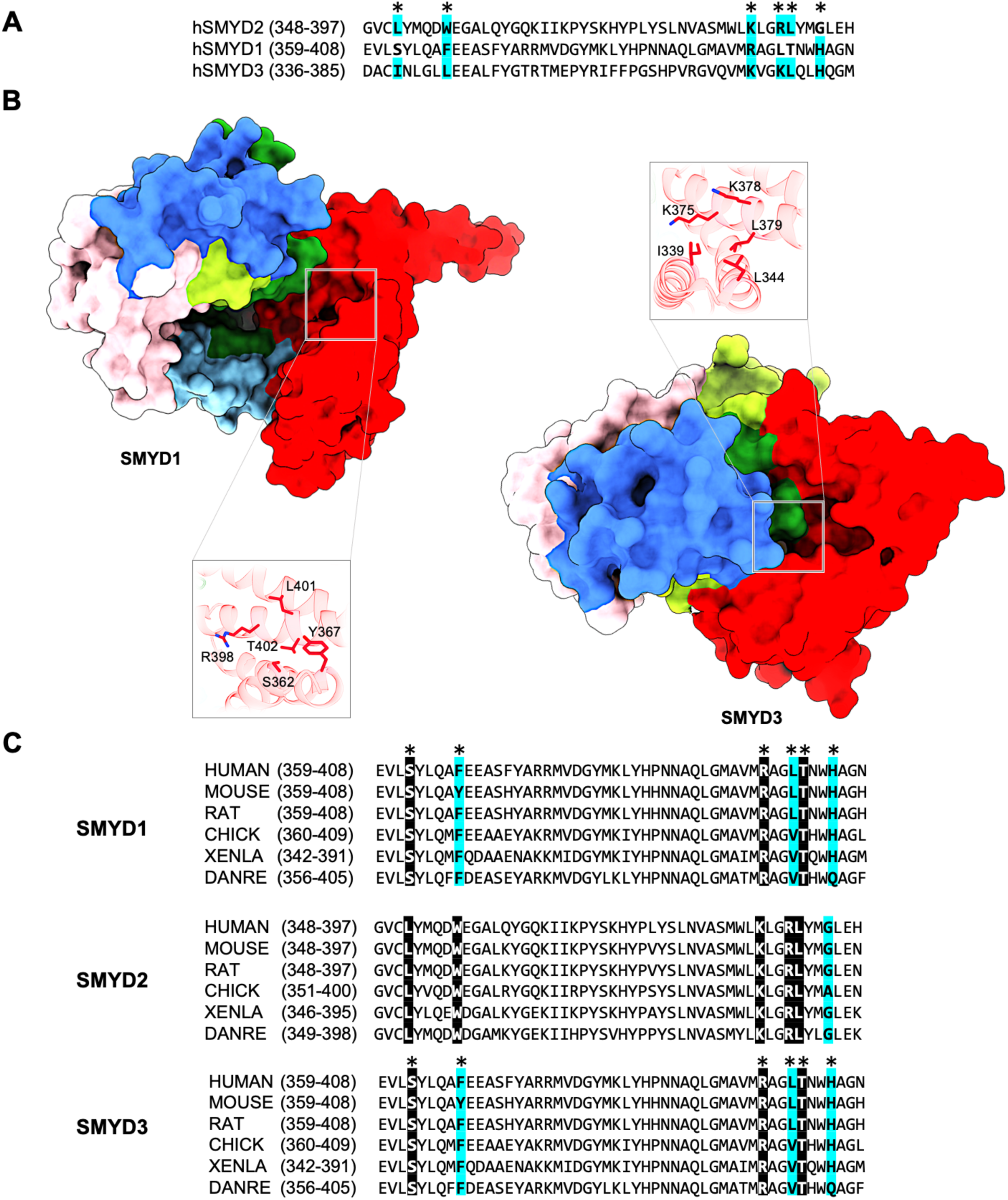
Putative secondary binding site in other SMYD proteins. (A) Sequence alignment of SMYD2, SMYD1, and SMYD3 at the secondary binding site. SMYD2 residues directly interacting with PARP1 are indicated by asterisks above the alignment. Completely identical residues are shown as white on black, and similar residues appear shaded in cyan. (B) A putative binding site in SMYD1 and SMYD3 at a location structurally equivalent to the secondary binding site in SMYD2. (C) Sequence alignment of SMYD proteins at the secondary binding site across species.

Given the extensive involvement of SMYD2 in various cancers and its demonstrated therapeutic value in preclinical studies (*17*), the discovery of this novel allosteric site opens exciting possibilities for cancer drug development. Targeting allosteric sites offers numerous advantages (*27*). They are often less conserved than active sites, allowing for increased selectivity and specificity for protein isoforms, which reduces off-target effects. This is crucial for SMYD2, as the human genome contains over 50 SET domain-containing methyltransferases with highly conserved active site structures, making off-target effects a concern (*28*). Allosteric modulators also tend to be safer since they do not completely shut down essential enzymatic activities, reducing toxicity. This feature is also especially important for SMYD2, given its ubiquitous expression in normal adult tissues, critical for fundamental cellular functions, including cell differentiation and proliferation, DNA repairing processes, and multiple signaling pathways (*12*). Our detailed study of this allosteric site lays a strong structural and biochemical foundation for future drug design. The site has an ideal mix of features for drug development: hydrophobic regions for strong binding, hydrophilic regions for solubility and specificity, and a moderately recessed pocket for stable drug access. These characteristics ensure high binding affinity, specificity, and optimal drug-like properties.

In conclusion, the discovery of this novel allosteric site not only reveals a new biochemical mechanism for SMYD2 regulation but also opens new therapeutic avenues for targeting SMYD2 in cancer. This allosteric site communicates structural and functional changes to the active site, while the active site does not influence the allosteric site. This unidirectional communication ensures that inhibitory signals received at the allosteric site can inhibit SMYD2 activity without being affected by substrate binding or product formation at the active site, offering robust inhibition.

## Materials and Methods

### Molecular cloning and mutagenesis

Full-length human SMYD2 (residues 1–433) (SMYD2-WT) and human PARP1 (residues 518–1014) were constructed by subcloning their DNA fragments into the pCDF-SUMO vector containing an N-terminal 6xHis-SUMO tag (*29*). SMYD2-F184A and SMYD2-L351A/W356A mutants were prepared using the Phusion Site-directed Mutagenesis Kit (New England Biolabs) according to the manufacturer’s instructions (*29*). GST-tagged SMYD2 proteins, including GST-SMYD2-WT, GST-SMYD2-F184A, and GST-SMYD2-L351A/W356A, were constructed by subcloning into the pGEX-6p-2 vector. The corresponding primers used in molecular cloning are listed in Table S1.

### Protein expression and purification

Proteins with the 6xHis-SUMO tag, including SMYD2-WT, SMYD2-F184A, SMYD2-L351A/W356A, and PARP1(518–1014), were expressed and purified according to previously described procedures (*19*). The 6xHis-SUMO tag was cleaved off with yeast SUMO Protease 1, generating a native N-terminus. Proteins with the GST tag, including GST alone, GST-SMYD2-WT, GST-SMYD2-F184A, and GST-SMYD2-L351A/W356A, were purified by glutathione sepharose affinity chromatography (*30*). Histone H3 and H4 were purified from inclusion bodies under denaturing conditions as previously described (*31*). Briefly, BL21 cells expressing histones were lysed by a French Press. Inclusion bodies were isolated by centrifugation and resuspended in denaturing buffer (7 M urea, 20 mM NaCl). Histones were purified from the inclusion bodies by cation-exchange chromatography and then refolded by dialysis against refolding buffer (10 mM NaCl, 1 mM β-mercaptoethanol).

### Crystallization and data collection

Co-crystallization of wild-type or mutant SYMD2 with AdoHcy and PARP1 was performed using the hanging-drop vapor diffusion method at 20°C (*32*). For PARP1, a 13- residue synthetic peptide (RMKLTLKGGAAVD) corresponding to human PARP1 residues 522–534 (CPC Scientific) was used for co-crystallization. All SMYD2 proteins, including SMYD2-WT, SMYD2-F184A, and SMYD2-L351A/W356A, were crystallized under essentially the same condition, which included 20% PEG3350, 100 mM Tris pH 7.5, 5% ethanol, 1 mM PARP1 peptide, and 600 μM AdoHcy. X-ray data from single crystals were collected at beamline 21-ID-F at the Advanced Photon Source (Argonne, IL) and processed using XDS (*33*). SMYD2-WT and SMYD2-L351A/W356A crystals were indexed to the *C*2 space group, containing two molecules per asymmetric unit. SMYD2-PEG and SMYD2-F184A crystals belong to the tetragonal space group *I*4 and contain one molecule per asymmetric unit (Table S2).

### Structure determination and refinement

Structures of SMYD2 complexes, including SMYD2-PARP1, SMYD2-F184A, SMYD2-L351A/W356A, and SMYD2-PEG, were solved by molecular replacement using a human SMYD2 structure (PDB code: 4WUY) as a search model. Manual model building was carried out in COOT (*34*), and refinement was performed using PHENIX (*35*). To reduce the effect of model bias, iterative-build omit maps were used during model building and structure refinement. The final models were analyzed and validated with MolProbity (*36*).

### Isothermal titration calorimetry

ITC experiments were carried out with a NanoITC SV calorimeter (TA Instruments) at 27°C. Proteins and peptides were dissolved in the same buffer (25 mM Tris pH 7.5, 50 mM NaCl), and their concentrations were quantified using a Direct Detect (Millipore Sigma). The sample cell contained 0.95 mL of 40 μM SMYD2 solution, and the injection syringe contained 250 μL of 631 μM PARP1 peptide solution. All titration experiments were performed with 16 injections of 15 μL peptide solution at intervals of 200 seconds. Control experiments performed by injections of the buffer into the protein solution yielded insignificant heats of dilution. Data were processed and fitted using NanoAnalyze (TA Instruments) to one-site, two-site, or cooperative models.

### GST pulldown assay

GST pulldown experiments were performed to investigate the interaction between SMYD2 and PARP1. Purified PARP1 (518–1014) was incubated with GST alone or GST tagged SMYD2 proteins in the binding buffer (50 mM Tris–HCl, pH 8.5, 100 mM NaCl, 10 mM DTT, 0.1% Triton X-100). After being pulled down with the glutathione agarose beads, the interactions were analyzed by SDS-PAGE and Coomassie Blue staining.

### ELISA protein binding assay

ELISA solid-phase protein binding experiments were performed according to the previously described protocol (*22*). ELISA plates were prepared by coating PARP1, H3, or H4 onto the wells in a coating buffer containing 10 mM HEPES pH 7.5, 0.1 M KCl, and 3 mM MgCl_2_. After blocking the plates with 1% BSA, binding experiments were performed by adding different concentrations of SMYD2 proteins into the wells. For PARP1 peptide inhibition experiments, SMYD2 proteins were pre-incubated with the PARP1 peptide before being added to the wells. Bound SMYD2 proteins were probed with the anti-SMYD2 antibody (ab108217, Abcam), and the amount of binding was quantified using an HRP-conjugated secondary antibody (ab6721, Abcam) that generated a color signal from oxidizing the substrate 2,2′-azino-bis(3-ethylbenzthiazoline sulfonic acid). The absorbance of signals was measured at 420 nm using a microplate reader at 2- minute intervals for 15 minutes. The rate of change in absorbance obtained by linear regression was used to quantify the binding. IC_50_ and the concentration required for 50% saturation were obtained by fitting binding data using a four-parameter logistic regression model. For reciprocal binding experiments, SMYD2 proteins were coated onto the wells and bound PARP1 was detected by the anti-PARP1 antibody (ab191217, Abcam). All binding experiments were performed in triplicate.

### Antibody-based methyltransferase assay

An antibody-based methyltransferase assay was carried out similarly to a previously described method (*29*). Reactions were performed by incubating SMYD2 proteins with H3 or H4 in the presence of AdoMet at 30°C overnight in a reaction buffer containing 50 mM Tris pH 8.5, 25 mM NaCl, 5% glycerol, and 2 mM DTT. Lysine methylation was detected by Western blot analysis using antibodies against mono-, di-, and trimethylated lysine (ab23366 & ab76118, Abcam). Enzymatic activities were quantified based on chemiluminescence using a CCD gel imager.

### Steady-state enzyme kinetics

Steady-state enzyme kinetics was carried out using the MTase-Glo methyltransferase assay according to the manufacturer’s instructions (Promega). Reactions were set up by incubating 2 μM SMYD2 proteins with varying concentrations of PARP1 peptides for different periods in the presence of a constant concentration of AdoMet (50 μM). After incubating the reactions with the MTase-Glo reagent and MTase-Glo Detection Solution, the amount of produced AdoHcy was determined by measuring luminescent signals using a microplate reader. The reactions were quantified using an AdoHcy standard curve that correlates the luminescence intensity to the AdoHcy concentration. Enzyme kinetic constants (*K_M_*, *k_ca_*_t_, and Hill coefficient) were determined using initial velocity measurements at varying substrate concentrations by fitting them to a standard Michaelis-Menten kinetics model and the Hill equation.

### Statistical analysis

GST-pulldown, ELISA, and methyltransferase assay results were presented as mean ± standard error (SE). Statistical differences between means were analyzed by one-way ANOVA if more than two groups were compared, and post-hoc comparisons on each pair of means were calculated using Tukey Honest Significant Difference (HSD) tests (SPSS Statistics). For comparisons between two groups, a two-tailed Student’s *t*-test was used. In both cases, a *p*-value ≤ 0.05 was considered statistically significant.

## Funding

This work was supported by the National Institutes of Health grant number 1R01HL128647-01 (to CL) and its subaward from the Georgia State University SP00012594 (to ZY).

### Author contributions

Y.Z., N.S., E.P. and Z.Y. designed the study; Y.Z., E.A., J.S., N.S., E.P. and J.B. performed the experiments. Y.Z., E.A., J.S., N.S., E.P. and J.B. collected and analyzed data; T.C., S.S., G.W., T.S., J.J., C.L. provided reagents; Y.Z. and Z.Y. wrote the manuscript.

### Data and materials availability

Coordinates and structure factors have been deposited in the Protein Data Bank (PDB) with accession numbers 9CKC for SMYD2-PARP1, 9CKF for SMYD2-L351A/W356A, 9CKG for SMYD2-F184A, and 6N3G for SMYD2-PEG. All other data needed to evaluate the conclusions in the paper are present in the paper and the Supplementary Materials.

