## Supplemental Materials for "Structure of the SMYD2-PARP1 Complex Reveals Both Productive and Allosteric Modes of Peptide Binding"

Yingxue Zhang *et al.*

**This PDF file includes:**

Supplementary Text
Figs. S1 to S3
Tables S1 to S2

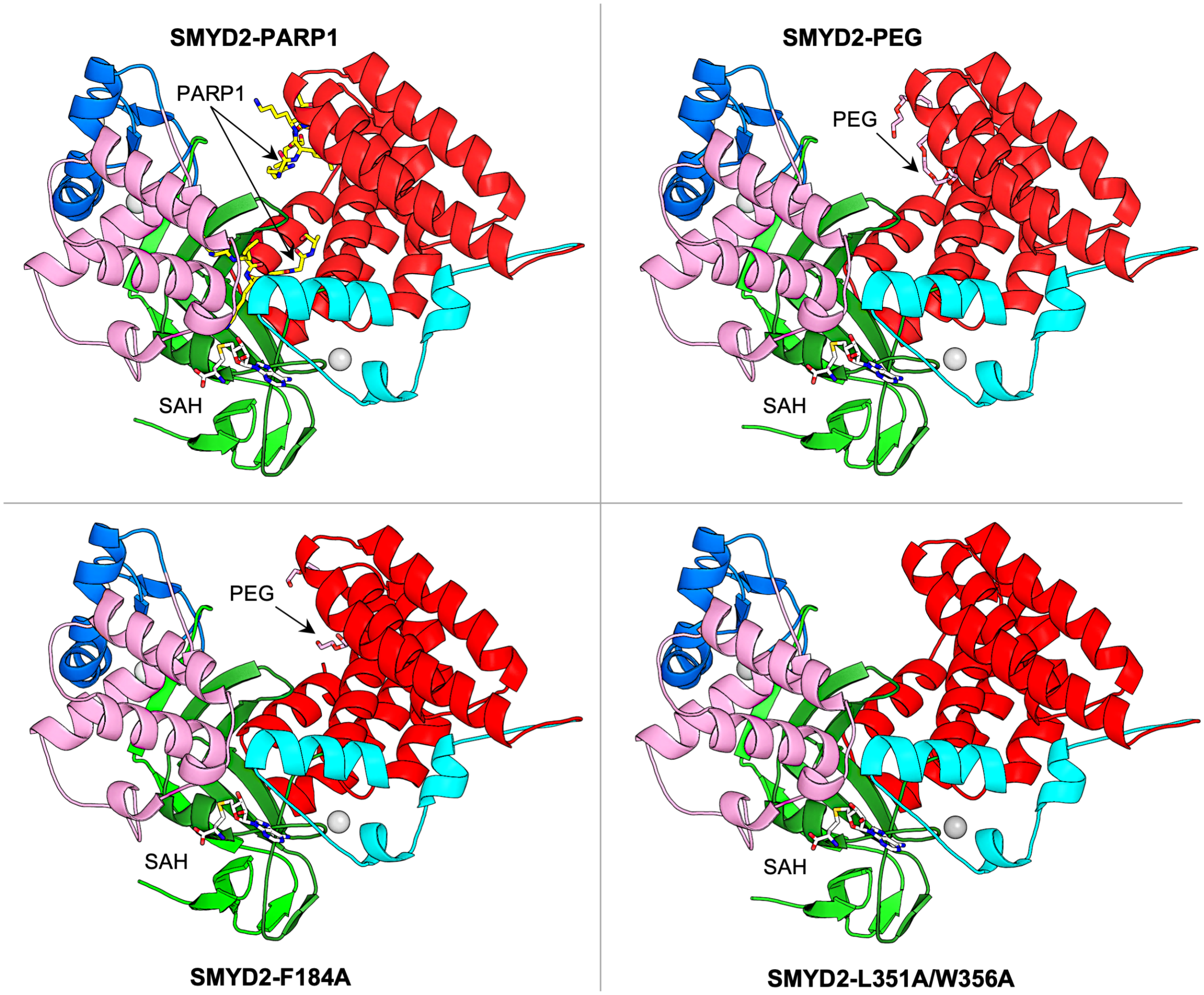


**Fig. S1. Overall structures of SMYD2-PARP1, SMYD2-PEG, SMYD2-F184A, and SMYD2-L351A/W356A.** The structures are colored according to domains, following the same scheme as in Figure 1A. AdoHcy, PARP1 peptides, and PEG are shown as sticks with carbon atoms colored in white, yellow, and pink, respectively. Zinc atoms are depicted as spheres.

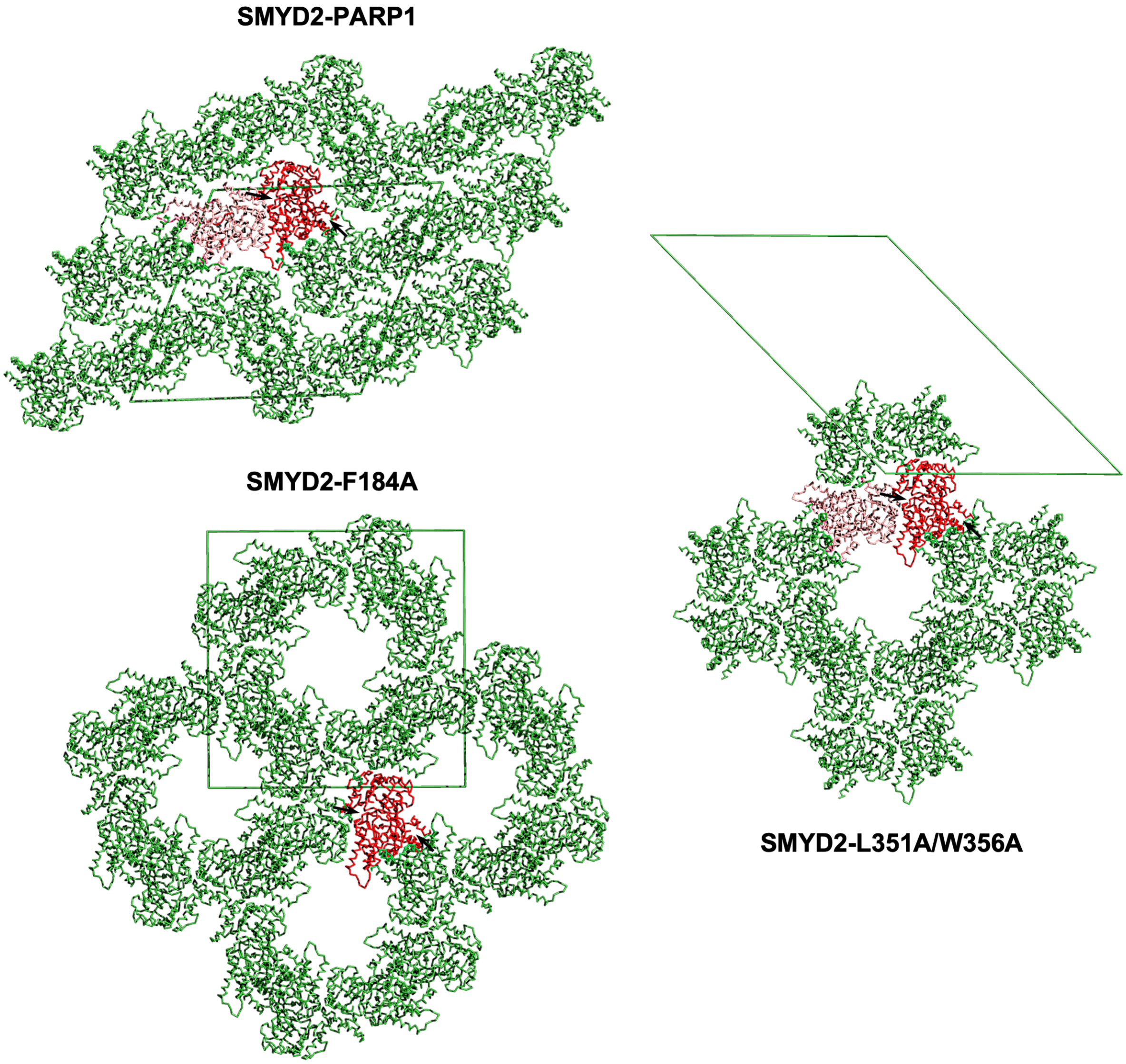


**Fig. S2. Crystal packing of the C2 and I4 forms.** SMYD2 is shown as a $C_\alpha$ trace, with red representing reference molecules and green representing symmetry-related molecules. The unit cell is shown as a green box. Left arrows point to the substrate binding site and right arrows to the secondary binding site.

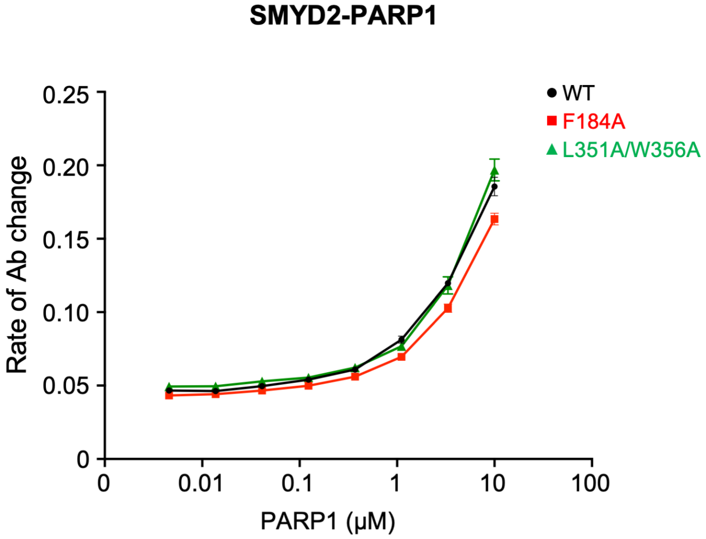


**Fig. S3. Reciprocal ELISA analysis of SMYD2-PARP1(518–1014) interaction.**

**Table S1. Molecular cloning primers**

| Construct | | Primer |
|---|---|---|
| SMYD2-PARP1 | F | 5′-GATCAAGGTCTCAAGGTATGAGGGCCGAGG-3' |
| | R | 5′-GCAGATCTCGAGTTATCAGTGGCTTTCAATTTCC-3' |
| SMYD2-F184A | F | 5′p-GGTTAACTGTAATGGCGCTACAATTGAAGATGAAG-3' |
| | R | 5′p-TGTGCAAAGAGTACTACGAGGC-3' |
| SMYD2-L351A/W356A | F | 5′p-CAGGACGCGGAAGGAGCCCTGCAATATGGACAGAAAATC-3' |
| | R | 5′p-CATGTACGCGCAGACACCCATGGCCTGGTACATCAT-3' |
| PARP1(518–1014) | F | 5′-TTATATCACCTGCATTAAGGTAAATCTGAAAAGAGAATG-3' |
| | R | 5′-ATCTATCTCGAGTTACCACAGGGAGGTCTTAAAATTGAA-3' |
| GST-SMYD2-WT | F | 5′-GTAATAGGATCCATGAGGGCCGAGGGC-3' |
| GST-SMYD2-F184A | R | 5′-GCAGATCTCGAGTTATCAGTGGCTTTCAATTTCC-3' |
| GST-SMYD2-L351A/W356A | | |

F: forward; R: reverse; p: phosphorylated primers

**Table S2. Crystallographic data and refinement statistics**

| | SMYD2-PARP1 | SMYD2-L351A/W356A | SMYD2-F184A | SMYD2-PEG |
|---|---|---|---|---|
| **Data** | | | | |
| Space group | *C*2 | *C*2 | *I*4 | *I*4 |
| Cell parameters | | | | |
| a, b, c (Å) | 142.8, 52.2, 144.9 | 215.2, 52.9, 151.9 | 151.8, 151.8, 53.4 | 151.8, 151.8, 53.7 |
| β (°) | 113.2 | 134.8 | | |
| Wavelength (Å) | 0.97856 | 1.07822 | 1.07822 | 0.97856 |
| Resolution (Å) | 133.2-2.1 (2.14-2.10)[a] | 107.89-2.50 (2.58-2.5)[a] | 75.9-2.75 (3.16-2.75)[a] | 107.4-2.43 (2.7-2.43)[a] |
| $R_{merge}$[b] | 0.105 (0.923) | 0.150 (1.225) | 0.121 (1.013) | 0.348 (1.927) |
| Redundancy | 7.3 (7.6) | 2.9 (2.9) | 6.1 (5.8) | 14.2 (13.8) |
| Unique reflections | 51097 | 27973 | 16090 | 14389 |
| Completeness (%) | 88.5 (100.0) | 82.6 (88.4) | 89.9 (61.2) | 94.8 (72.2) |
| ⟨I/σ⟩ | 13.7 (2.2) | 7.3 (1.4) | 9.3 (1.7) | 10.3 (1.6) |
| **Refinement** | | | | |
| Resolution (Å) | 66.6-2.1 (2.14-2.10) | 53.94-2.50 (2.60-2.50) | 48.0-2.75 (2.92-2.75) | 75.92-2.43 (2.62-2.43) |
| Molecules/AU | 2 | 2 | 1 | 1 |
| $R_{work}$[c] | 0.184 (0.265) | 0.222 (0.297) | 0.194 (0.398) | 0.175 (0.309) |
| $R_{free}$[d] | 0.208 (0.267) | 0.249 (0.330) | 0.223 (0.521) | 0.222 (0.373) |
| Ramachandran plot | | | | |
| Residues in favored | 97.5% | 96.7% | 98.4% | 92.3% |
| Residues in allowed | 2.5% | 3.3% | 1.6% | 6.3% |
| RMSD | | | | |
| Bond lengths (Å) | 0.004 | 0.003 | 0.006 | 0.011 |
| Bond angels (°) | 0.718 | 0.671 | 1.075 | 1.313 |
| No. of atoms | | | | |
| Protein | 6930 | 6935 | 3432 | 3462 |
| Peptide | 152 | - | - | - |
| Water | 420 | 125 | 0 | 79 |
| B-factor (Å$^2$) | | | | |
| Protein | 47.4 | 54.0 | 84.1 | 38.44 |
| Peptide | 71.5 | - | - | - |
| Water | 46.2 | 41.0 | - | 32.73 |

[a]Numbers in parentheses refer to the highest resolution shell.
[b]$R_{merge}$= Σ|I-⟨I⟩|/ΣI, where I is the observed intensity and ⟨I⟩ is the averaged intensity of multiple observations of symmetry-related reflections.
[c]$R_{work}$= $\Sigma|F_o-F_c|/\Sigma|F_o|$, where $F_o$ is the observed structure factor, $F_c$ is the calculated structure factor.
[d]$R_{free}$ was calculated using a subset (5%) of the reflection not used in the refinement.
